# Cardiomyocytes Undergo a Mesenchymal-Like Fate Transition in Myocardial Fibrosis

**DOI:** 10.64898/2026.06.10.731493

**Authors:** Tianle Wang, Chunyu Zhou, Mengdi Liu, Yujia Xing, Chunguang Han, Runhan Li, Yu Huang, Zhenhua Li, Yan Teng, Guan Yang, Wenjia Liu, Ping Xu, Shiqiang Wang, Bin Zhou, Jing-Dong J. Han, Jian Wang, Xiao Yang

## Abstract

**BACKGROUND:** Myocardial fibrosis, a pathological hallmark of adverse cardiac remodeling and heart failure, has been conventionally attributed to the activation of resident fibroblasts. Although recent studies suggest contributions from non-fibroblast lineages, direct *in vivo* genetic evidence that cardiomyocytes can undergo a mesenchymal-like fate transition during myocardial fibrosis remains absent. This study aims to investigate whether such a transition occurs and to elucidate the underlying regulatory mechanisms.

**METHODS:** Human myocardial infarction (MI) tissues were analyzed by immunohistochemistry and integrated with public single-nucleus RNA sequencing (snRNA-seq) data to detect mesenchymal-like signatures in cardiomyocytes. Genetic lineage-tracing was performed in MI mice, and in cardiomyocyte-specific *Hgs* (hepatocyte growth factor-regulated tyrosine kinase substrate) gene knockout mice, to map the fate of cardiomyocyte-derived cells. Mechanistic insights were obtained through proteomic and snRNA-seq analysis of *Hgs* knockout hearts and validated through gain- and loss-of-function experiments targeting *Aldh1a2* (aldehyde dehydrogenase 1 family member A2).

**RESULTS:** In human MI samples, a subset of cardiomyocytes showed reduced expression of cardiomyocyte markers concurrent with acquisition of mesenchymal-associated markers. Genetic lineage tracing demonstrated that adult cardiomyocytes can adopt a mesenchymal-like cell fate during post-MI remodeling. We identify HGS as a factor constraining this transition. *Hgs* knockout in adult cardiomyocytes upregulated *Aldh1a2,* triggered the mesenchymal-like fate transition, and gave rise to cells expressing markers of activated fibroblasts or osteoblasts, accompanied by pronounced myocardial fibrosis and calcification. Forced *Aldh1a2* overexpression in cardiomyocytes drove the mesenchymal-like fate transition *in vitro* and *in vivo,* whereas *Aldh1a2* deletion in cardiomyocytes mitigated MI-induced myocardial fibrosis.

**CONCLUSIONS:** This study provides *in vivo* genetic evidence that adult cardiomyocytes possess the capacity to undergo a mesenchymal-like fate transition under pathological conditions. Our data suggest that HGS and ALDH1A2 serve as regulators of the transition, offering a new basis for understanding cellular and molecular mechanisms of myocardial fibrosis.

**Novelty and Significance:** *What Is Known?:*

- Myocardial fibrosis is primarily driven by resident fibroblast activation, with additional contributions from cardiac CD34^+^ cells, pericytes, and macrophages.
- Adult cardiomyocytes exhibit phenotypic plasticity and transdifferentiate into epicardial-like or pacemaker cells under specific conditions.

*What New Information Does This Article Contribute?:*

- A subset of cardiomyocytes adopts a mesenchymal-like cell fate during myocardial fibrosis, marked by downregulation of cardiomyocyte identity markers and loss of aligned cell–cell contacts.
- These cells acquire mesenchymal morphology and markers, ECM components, migratory gene signatures, and proliferative capacity.
- HGS and ALDH1A2 act as regulators of this mesenchymal-like fate transition. Myocardial fibrosis drives heart failure progression, yet the cellular sources of pathological fibroblasts remain incompletely defined. Here, we demonstrate that a subset of cardiomyocytes adopts a mesenchymal-like cell fate during myocardial fibrosis by using an integrated approach combining human MI samples, murine genetic lineage tracing, and snRNA-seq. Mechanistically, we identify HGS and ALDH1A2 as regulators of this transition. Cardiomyocyte-specific *Hgs* deletion upregulates *Aldh1a2*, triggering the mesenchymal-like fate transition. Furthermore, *Aldh1a2* overexpression drives this transition, while its deletion attenuates MI-induced fibrosis. These findings reveal a previously unrecognized plasticity of adult cardiomyocytes and identify potential therapeutic targets for fibrotic heart disease.

## INTRODUCTION

Myocardial fibrosis, a maladaptive process characterized by excessive deposition of fibrous extracellular matrix (ECM) components in the cardiac interstitium, represents a convergent pathological hallmark of diverse cardiac diseases and a major driver of heart failure.^1,2^ Despite its clinical relevance, the underlying cellular mechanisms remain incompletely understood.

Cardiac fibroblasts are highly heterogeneous and play a pivotal role in myocardial fibrosis, serving dual functions in cardiac homeostasis and pathological remodeling.^3–6^ Under physiological conditions, these cells maintain myocardial structural integrity and ECM homeostasis. In response to injury, they undergo phenotypic switching to acquire a pro-fibrotic state, driven by a complex network of signaling pathways, including transforming growth factor-β (TGF-β), platelet-derived growth factor (PDGF), WNT signaling, and inflammatory cytokines.^7–10^ Activated cardiac fibroblasts differentiate into myofibroblasts that express contractile proteins such as α-smooth muscle actin (α-SMA) and matricellular protein periostin (POSTN), driving aberrant collagen deposition.^11,12^ Notably, cardiac fibroblasts can also transdifferentiate into osteoblasts, contributing to myocardial calcification after injury.^13^

Although resident fibroblast activation dominates myocardial fibrogenesis,^14–17^ lineage tracing studies reveal additional cellular sources, including epicardial-derived cells via epithelial-mesenchymal transition (EMT) or endothelial cells via endothelial-mesenchymal transition (EndMT),^18–20^ and transdifferentiation of cardiac CD34^+^ cells, pericytes, and macrophages under conditions such as pressure overload, diabetic cardiomyopathy, or myocardial infarction (MI).^21–23^ Whether cardiomyocytes can contribute to the fibroblast pool through a mesenchymal-like fate transition remains unclear.

Once considered terminally differentiated, cardiomyocytes are now recognized to retain plasticity under certain conditions.^24^ During mammalian cardiac regeneration following MI or other injury, adult cardiomyocytes undergo dedifferentiation, proliferation, and then redifferentiation into new cardiomyocytes via Hippo-YAP inhibition, ERBB2 activation, non-coding RNA regulation, or metabolic reprogramming.^25–34^ Partial reprogramming via OSKM (Oct4, Sox2, Klf4, c-MYC) factors in mice can transiently revert cardiomyocytes to a fetal state, while prolonged induction leads to irreversible dedifferentiation and risks tumorigenesis.^35^ Moreover, an intermediate cardiac reprogramming stage during atrial-to-ventricular cardiomyocyte transdifferentiation contributes to zebrafish cardiac ventricular regeneration.^36^

Emerging evidence also suggests that cardiomyocytes can reprogram into cardiac non-myocytes. Recent single-cell RNA sequencing studies have detected endothelial or fibroblast-like gene expression signatures in cardiomyocytes under conditions of pressure overload and atrial fibrillation.^37–40^ Ectopic Wt1 expression in zebrafish cardiomyocytes can induce epicardial-like cell conversion,^41^ while Tbx18 expression or biomaterials treatment in rodent cardiomyocytes can directly reprogram them into pacemaker cells.^42,43^ Additionally, ERBB2 overexpression in postnatal day 7 (P7) mouse cardiomyocytes drives EMT-like processes during cardiac regeneration.^44^ These findings collectively challenge the conventional understanding of cardiomyocyte lineage restriction, yet direct *in vivo* evidence demonstrating a mesenchymal-like fate transition of adult cardiomyocytes in myocardial fibrosis remains limited.

In this study, we investigate a previously underexplored aspect of adult cardiomyocyte plasticity in the context of myocardial fibrosis. By integrating lineage tracing and single-nucleus RNA sequencing (snRNA-seq), we provide *in vivo* evidence that a subset of cardiomyocytes undergoes a mesenchymal-like fate transition in the setting of myocardial fibrosis. Under conditions of cardiac injury and genetic perturbation, these cardiomyocytes downregulate cardiomyocyte identity markers, acquire mesenchymal features and a spindle-shaped morphology, and give rise to cells expressing myofibroblast or osteoblast markers. These findings raise the possibility that cardiomyocytes may represent one potential cellular source contributing to the fibroblast population in myocardial fibrosis.

## METHODS

### Data Availability

The authors declare that all supporting data are available in the article and its online supplementary files, and are available from the corresponding authors upon reasonable request.

### Human Subjects

Analyses involving human samples followed the principles outlined in the Declaration of Helsinki. Detailed protocols and subject information are provided in the Supplemental Material.

### Animal Models

All animal experimental protocols were approved by the institutional animal care and use committee of the National Center for Protein Sciences (Beijing) (approval number NCPSB-20240222-18MBL) and conformed to the US National Institutes of Health Guide for the Care and Use of Laboratory Animals. Extended details of the animal models, anesthesia, and follow-up studies are described in the Supplemental Material.

### Quantitative and Statistical Analyses

For simple two–group comparison, normality was assessed using the Shapiro–Wilk test and equality of variances using the *F*-test. When these assumptions were met, unpaired two-tailed Student’s *t* tests were applied. Two-way ANOVA followed by Tukey’s multiple comparisons test was used to analyze control and *α-MHC-Cre*;*Aldh1a2^fl/fl^* mice subjected to sham or MI. Non-normally distributed data were analyzed using the Mann-Whitney *U* test. All data are presented as mean ± SEM and were derived from at least 3 independent experiments. Results with *p* < 0.05 were considered statistically significant.

Throughout the study, *n* denotes independent biological replicates (individual mice or human donor hearts), unless otherwise indicated. The number of replicates (n) is indicated in figure legends. For RNA–seq analysis, 3 independent pools were generated per group, each generated by pooling 3–4 mice. For snRNA–seq, samples were prepared by pooling 2 mice (*Hgs^+/+^*) or 4 mice (*Hgs^fl/fl^*).

To quantify α-SMA-positive cells expressing EYFP or EYFP-labeled cardiomyocytes expressing α-SMA in the hearts of sham and MI mice, six groups of mice were analyzed. For each heart, 7–8 regions containing positive signals were assessed.

## RESULTS

### Cardiomyocytes Show a Mesenchymal-Like Transition in Human Infarcted Myocardium

To investigate the potential plasticity of cardiomyocytes in MI, we analyzed left ventricular cardiomyocyte nuclei from both the lesion and border regions of 17 MI patients, as well as from 4 non-transplanted donor hearts, utilizing publicly available snRNA-seq dataset.^45^ A total of 51,930 nuclei were profiled and identified as cardiomyocytes based on the expression of canonical markers, including *FHL2*, *TTN*, *RYR2*, *MYH7*, and *MYL2* (Figure 1A). In MI patients, cardiomyocytes showed a decrease in cardiomyocyte identity markers and an increase in mesenchymal transcriptional factors (*TWIST1, SNAIL1* and *SNAIL2*) and mesenchymal markers (*PDGFRα* and *PDGFRβ*). These transcriptional alterations suggest a possible mesenchymal-like transition during MI (Figure 1B).

**Figure 1.**
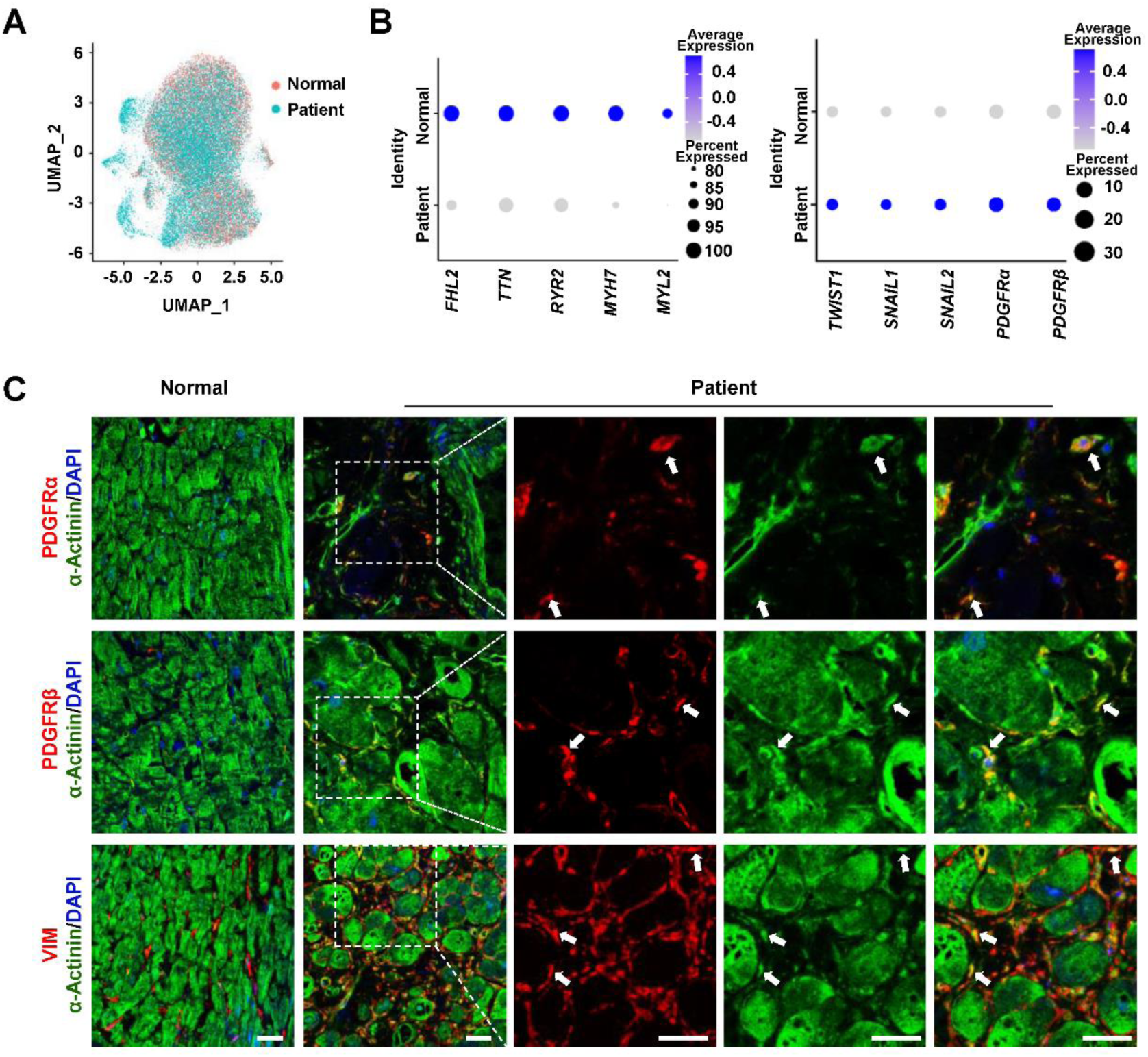
A subset of cardiomyocytes exhibits mesenchymal-like characteristics in the hearts of MI patients. **A**, UMAP visualization of cardiomyocyte clusters from normal human and MI patients. **B**, Dot plots showing expression levels of representative cardiomyocyte and mesenchymal cell markers in cardiomyocytes from normal human and MI patients. Color intensity indicates expression levels. Dot size reflects the percentage of cells expressing the given genes. **C**, Representative immunofluorescence images showing co-localization of mesenchymal markers (red) with α-Actinin (green) in cardiac sections from normal human and MI patients. Scale bars, 25 μm.

To validate these observations, we performed immunofluorescence staining on heart sections from MI patients (Table S1) and controls. In normal human hearts, mesenchymal markers PDGFRα, PDGFRβ, and VIMENTIN (VIM) were confined to α-Actinin^-^ mesenchymal-like interstitial cells. In contrast, MI hearts showed co-localization of these mesenchymal markers with the cardiomyocyte marker α-Actinin (Figure 1C). Notably, many of these double-positive cells frequently displayed a spindle-shaped morphology distinct from the characteristic cylindrical, striated architecture of cardiomyocytes, resembling mesenchymal-like cells (Figure 1C). Z-stack confocal microscopy verified intracellular co-expression, ruling out signal overlap from adjacent cells (Figure S1).

Collectively, these data indicate that cardiomyocytes acquire molecular and morphological features indicative of a possible mesenchymal-like fate transition during MI.

### Cardiomyocytes Undergo a Mesenchymal-Like Fate Transition in MI Mice

To determine whether cardiomyocytes undergo a mesenchymal-like fate transition *in vivo*, we established an MI model using *α-MHC-MerCreMer*;*ROSA26^LSL-EYFP^* (hereafter *α-MHC-MCM*;*ROSA26^EYFP^*) mice for lineage tracing. Cre recombinase specificity was verified in 2-month-old *α-MHC-MCM;ROSA26^EYFP^* mice. Immunofluorescence analysis confirmed that Cre activity was restricted to cardiac troponin T^+^ (cTnT^+^) cardiomyocytes but not detected in CD31^+^ endothelial cells, α-SMA^+^ smooth muscle cells, Sca-1^+^ progenitor cells, PDGFRβ^+^ mural cells, or VIM^+^ fibroblasts (Figure S2A), consistent with previous reports.^46–48^ Following tamoxifen administration to label cardiomyocytes and their progeny with EYFP, a 21-day washout period was implemented to ensure complete tamoxifen clearance. Mice then underwent either sham surgery or permanent coronary artery ligation to induce MI, with cardiac tissues harvested 5 days post-operation (Figure 2A).

**Figure 2.**
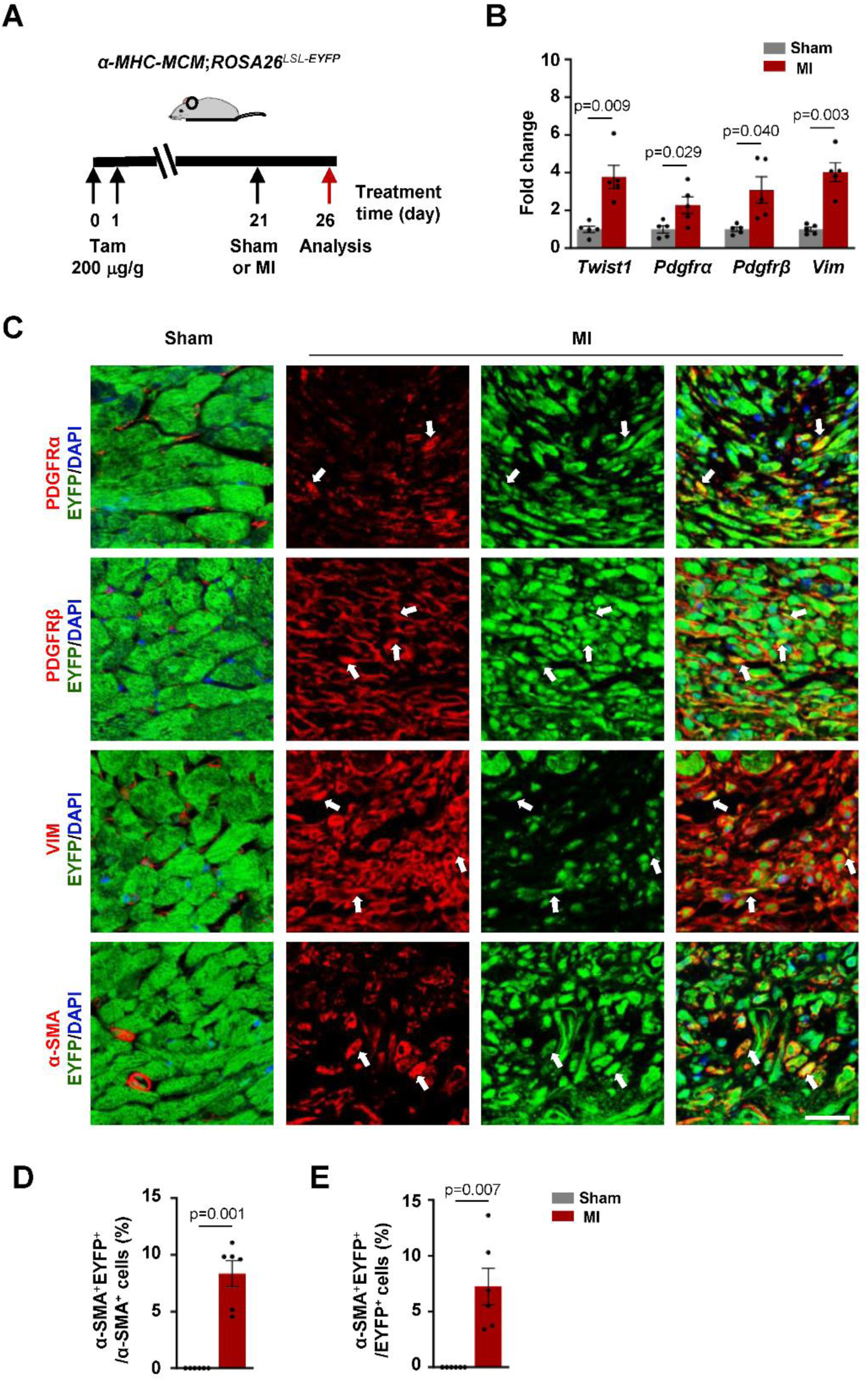
Cardiomyocytes undergo a mesenchymal-like fate transition in MI mice. **A**, Schematic diagram of lineage tracing in 2-month-old *α-MHC-MCM;ROSA26^EYFP^* mice. **B**, RT-qPCR analysis of mRNA levels of mesenchymal marker genes in adult cardiomyocytes isolated from sham and MI mice. **C**, Representative immunofluorescence images showing co-localization of mesenchymal markers (red) with EYFP (green) in cardiac sections from sham and MI mice. Scale bar, 25 μm. **D**, Quantification of α-SMA positive cells expressing EYFP in hearts of sham or MI mice. **E**, Quantification of EYFP-labeled cardiomyocytes expressing α-SMA in hearts of sham or MI mice. Data in **B**, **D** and **E** were presented as means ± SEM. *P* values were determined by unpaired two-tailed Student’s *t* test. Number of samples is shown on bar graphs.

Quantitative real-time PCR (RT-qPCR) analysis of isolated adult cardiomyocytes revealed significant upregulation of mesenchymal transcriptional factor *Twist1*, along with the mesenchymal-associated markers *Pdgfrα*, *Pdgfrβ*, and *Vim* in MI mice compared to sham controls (Figure 2B). Immunofluorescence staining further showed that some EYFP-labeled cardiomyocytes-derived cells within fibrotic foci co-expressed the mesenchymal markers PDGFRα, PDGFRβ, and VIM (Figure 2C). Additionally, a subset of EYFP^+^ cells expressed myofibroblast marker α-SMA in MI mice (Figure 2C). Quantitative analysis of the infarct and peri-infarct regions showed that approximately 8.3% ± 1.1% of α-SMA^+^ cells originated from cardiomyocytes, while 7.2% ± 1.6% of EYFP^+^ cardiomyocyte-derived cells acquired α-SMA expression (Figure 2D and 2E).

Morphologically, some EYFP-labeled cells in MI hearts exhibited disrupted cell-cell contacts, loss of directional alignment, and a shift toward a spindle–shaped appearance reminiscent of mesenchymal cells (Figure 2C). In contrast, hearts from sham-operated controls maintained characteristic organized, tightly connected cardiomyocyte architecture (Figure 2C).

Taken together, these lineage tracing data provide *in vivo* evidence that adult cardiomyocytes can undergo a mesenchymal-like fate transition following ischemic injury.

### *Hgs* Deletion in Cardiomyocytes Drives a Mesenchymal-Like Fate Transition

Gene Ontology (GO) and Reactome pathway analyses of human cardiomyocytes revealed that signaling pathways associated with endosomal transport, ubiquitination, autophagy, and receptor tyrosine kinases were significantly downregulated in MI patients (Figure S3A and 3B). Given that hepatocyte growth factor-regulated tyrosine kinase substrate (HGS) is a core component of the endosomal sorting complex required for transport (ESCRT) involved in regulating these processes, and based on our prior finding that cardiomyocyte-specific *Hgs* knockout induces myocardial fibrosis and calcification,^49^ we hypothesized that *Hgs* deficiency might drive cardiomyocytes to undergo a mesenchymal-like fate transition.

To test this possibility, we performed genetic lineage tracing utilizing *α-MHC-Cre*;*Hgs^fl/fl^*;*ROSA26^EYFP^* (hereafter *Hgs^fl/fl^*;*ROSA26^EYFP^*) mice, in which α-MHC-Cre recombinase activity is restricted to cardiomyocytes (Figure S2B). These mice were not subjected to MI surgery, and thus any observed phenotypes arise solely from the genetic manipulation. As expected, *Hgs* knockout hearts exhibited prominent myocardial fibrosis and calcification, confirmed by morphological assessments, Masson’s trichrome and Von Kossa staining of heart sections (Figure S4A). Flow cytometry identified a distinct population of EYFP^+^PDGFRα^+^ cells, comprising 0.97% ± 0.13% of viable cells from *Hgs* knockout hearts (Figure 3A). Immunofluorescence staining showed that a subset of lineage-traced EYFP^+^ cardiomyocyte-derived cells co-expressed the mesenchymal markers PDGFRα and VIM in *Hgs* knockout mice (Figure 3B; Figure S4B), which was further confirmed by Z-stack confocal images (Figure S4C). These cells displayed disrupted cell-cell contacts and reduced expression of the cardiomyocyte marker cTNT (Figure S4D).

**Figure 3.**
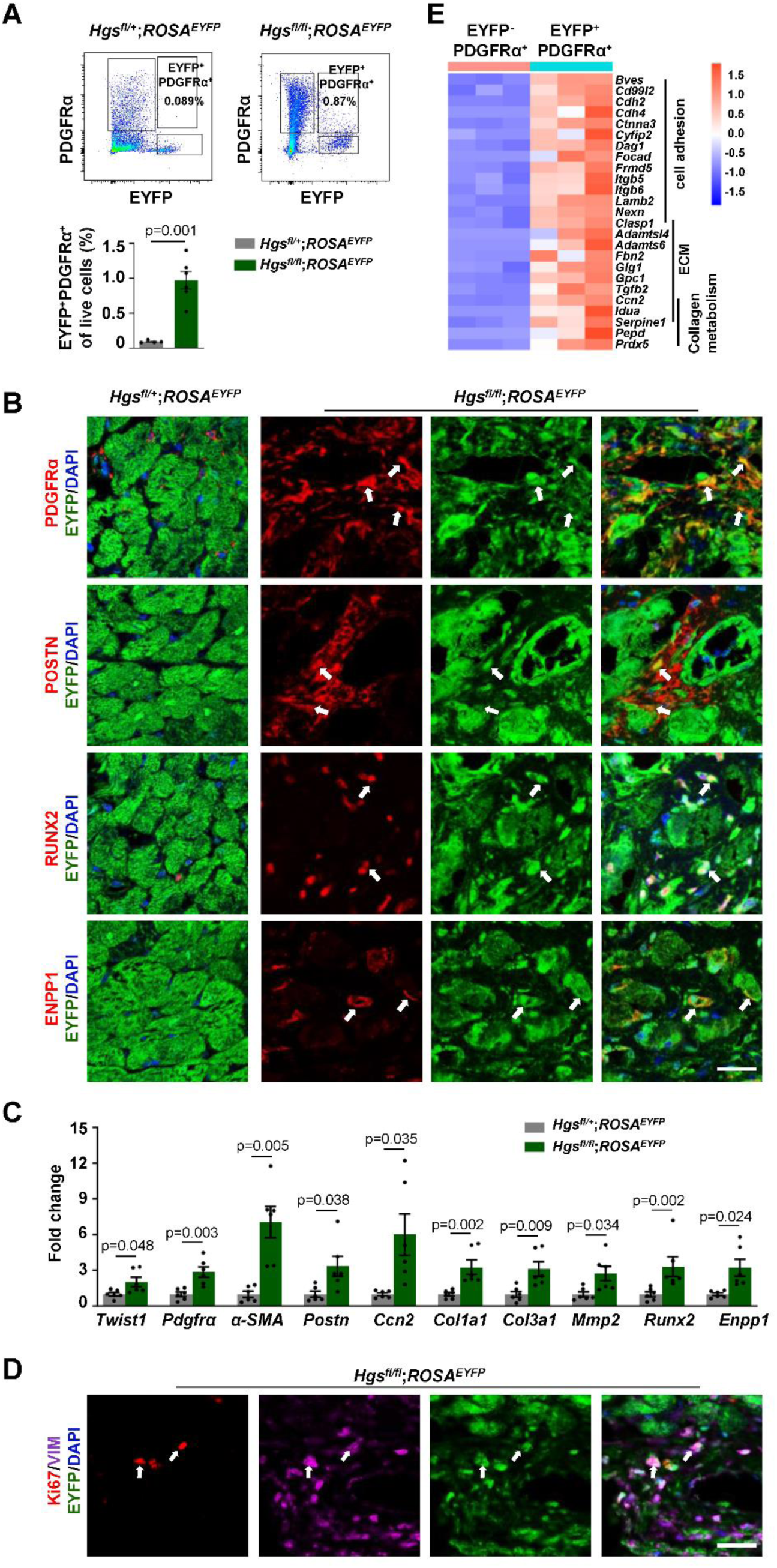
*Hgs* deletion in cardiomyocytes drives a mesenchymal-like fate transition. **A**, Flow cytometry analysis quantifying the EYFP^+^Pdgfrα^+^ cell population in cardiac tissues from mice. *Hgs^fl/+^;ROSA^EYFP^* refer to *α-MHC-Cre*;*Hgs^fl/+^*;*ROSA^EYFP^*, *Hgs^fl/fl^;ROSA^EYFP^* refer to *α-MHC-Cre*;*Hgs^fl/fl^*;*ROSA^EYFP^*. Bar graph shows the percentage of EYFP^+^Pdgfrα^+^ cells among live cells. **B**, Representative immunofluorescence images showing co-expression of mesenchymal cell, activated fibroblast, and osteoblast markers (red) with EYFP (green) in cardiac sections from mice. Scale bar, 25 μm. **C**, RT-qPCR analysis of mRNA levels of mesenchymal transcription factor, mesenchymal marker genes, ECM-associated molecules and osteoblast markers in adult cardiomyocytes isolated from mice. **D**, Representative immunofluorescence images showing co-expression of Ki67 (red) and VIM (purple) with EYFP (green) in cardiac sections from *Hgs^fl/fl^*;*ROSA^EYFP^* mice. Scale bar, 25 μm. **E**, Clustering heatmap of differentially expressed genes associated with cell adhesion, ECM and collagen in EYFP^+^Pdgfrα^+^ versus EYFP^-^Pdgfrα^+^ cells from *Hgs^fl/fl^*;*ROSA^EYFP^* mice. p < 0.05, log2(FC)>1. Data are from 3 biological replicates per group (3–4 mice pooled per replicate). Data in **A** and **C** were presented as means ± SEM. *P* values were determined by unpaired two-tailed Student’s *t* test (normally distributed data) or Mann-Whitney *U* test (non-normally distributed data). Number of samples is shown on bar graphs.

Given the prominent fibrosis and calcification in *Hgs* knockout hearts, we examined whether cardiomyocyte-derived mesenchymal-like cells further acquired myofibroblast and osteoblast features. In *Hgs* knockout mice, a subset of EYFP^+^ cells co-expressed myofibroblast markers α-SMA and POSTN, osteogenic markers RUNX2 and OPN, and ENPP1—a key enzyme in bone mineralization (Figure 3B; Figure S4B). RT-qPCR analysis confirmed coordinated upregulation of mesenchymal transcription factor *Twist1*, mesenchymal marker *Pdgfrα*, myofibroblast marker gene *α-SMA*, ECM genes (*Ccn2*, *Col1a1*, *Col3a1*), migration associated ECM components (*Postn* and *Mmp2*), and osteogenic markers (*Runx2* and *Enpp1*) in adult cardiomyocytes isolated from *Hgs* knockout mice (Figure 3C). Additionally, a proportion of cardiomyocyte-derived mesenchymal-like cells were proliferative, as demonstrated by co-staining for EYFP, VIM, and Ki67 (Figure 3D).

To characterize the transcriptional profile of cardiomyocyte-derived mesenchymal-like cells relative to mesenchymal cells from other sources, we performed fluorescence-activated cell sorting (FACS) to isolate EYFP^+^PDGFRα^+^ (cardiomyocyte-derived) and EYFP^-^PDGFRα^+^ (non-cardiomyocyte-derived) populations from *Hgs* knockout mice for transcriptomic analysis. Cardiomyocyte-derived mesenchymal-like cells showed elevated expression of genes associated with cell adhesion, ECM organization, collagen production, migration, and proliferation (Figure 3E; Figure S4E). Moreover, compared to mesenchymal cells from other origins, the cardiomyocyte-derived subset displayed distinct transcriptional signatures related to cell morphology, fate determination, differentiation, and osteogenic/bone-associated genes (Figure S4F and S4G).

We further validated these findings using an inducible cardiomyocyte-specific *Hgs* knockout mouse model with an EYFP reporter gene (*α-MHC-MCM*;*Hgs^fl/fl^;ROSA26^EYFP^*, hereinafter *iHgs^fl/fl^;ROSA26^EYFP^*). Two months after tamoxifen administration (200 µg/g daily for five days) (Figure S5A), *iHgs^fl/fl^;ROSA26^EYFP^* mice recapitulated the fibrocalcific phenotype and exhibited similar molecular alterations (Figure S5B and S5C). Cardiomyocytes isolated from these mice showed significant upregulation of the mesenchymal transcriptional factor *Twist1*, mesenchymal markers (*Pdgfrα*, *Vim*), myofibroblast marker *α-SMA*, ECM genes (*Fn1*, *Ddr2*, *Col1a1*, and *Col3a1*), and migration associated ECM components (*Postn*, *MMP2*) (Figure S5B). Furthermore, a subset of EYFP-labeled cardiomyocytes-derived cells in *iHgs^fl/fl^;ROSA26^EYFP^* mice co-expressed PDGFRα, VIM, α-SMA, RUNX2, OPN, and OCN (Figure S5C).

Together, these results suggest that *Hgs* deficiency in cardiomyocytes activates a mesenchymal–like program, characterized by upregulation of mesenchymal identity genes, ECM and migration factors, and acquisition of proliferative capacity, alongside partial loss of cardiomyocyte identity. This provides *in vivo* evidence that cardiomyocytes can undergo a mesenchymal-like fate transition in the context of myocardial fibrosis caused by genetic disturbance.

#### Cardiomyocyte-Derived Mesenchymal-Like Cells Give Rise to Cells Expressing Myofibroblast or Osteoblast Markers in *Hgs* Knockout Mice

To unambiguously track the fate of cardiomyocyte-derived mesenchymal-like cells, we performed dual recombinase-based genetic lineage tracing by generating an *α-MHC-Cre;Pdgfrβ^LSL-Dre^;Hgs^fl/fl^;ROSA26^LSL-EYFP/RSR-tdTomato^* mouse line. In this system, *Pdgfrβ^LSL-Dr^*^e^ allele contains a *loxP*-flanked STOP cassette preceding the Pdgfrβ-Dre (Figure 4A). α-MHC-Cre activity leads to permanent EYFP labeling of cardiomyocytes and their progeny. Concurrently, Cre-mediated excision of the STOP cassette enables Dre expression under the control of the Pdgfrβ promoter. As a result, only cardiomyocyte-derived cells that subsequently express Pdgfrβ will be labeled with both EYFP and tdTomato (Figure 4A).

**Figure 4.**
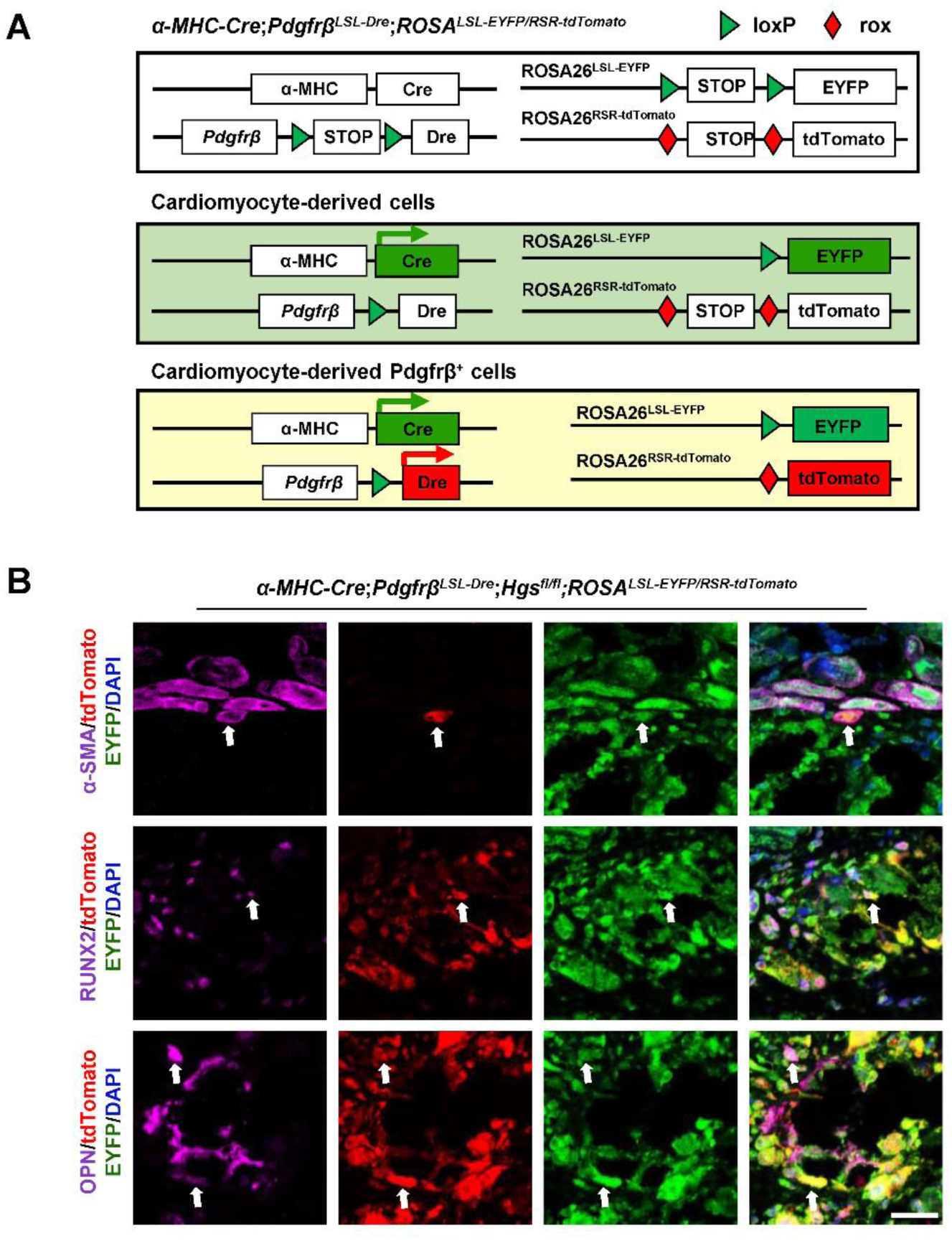
Cardiomyocyte-derived mesenchymal-like cells give rise to cells expressing myofibroblast or osteoblast markers in *Hgs* knockout mice. A, Schematic depicting dual recombinase-based genetic lineage using the *α-MHC-Cre*;*Pdgfrβ^LSL-Dre^*;*Hgs^fl/fl^*;*ROSA26^LSL-EYFP/RSR-tdTomato^* mouse line. **B**, Representative immunofluorescence images showing co-expression of myofibroblast/osteoblast markers (purple), tdTomato (red) with EYFP (green) in heart sections from mice. Scale bar, 25 μm.

In *Hgs* knockout mice, we detected a population of EYFP^+^tdTomato^+^ cells (Figure 4B). These double-positive cells co-expressed the myofibroblast marker α-SMA, the osteogenic transcription factor RUNX2, or the bone matrix protein OPN (Figure 4B). Taken together, these data provide genetic evidence consistent with the interpretation that, in the setting of *Hgs* deletion, cardiomyocyte-derived mesenchymal-like cells can subsequently give rise to cells expressing myofibroblast or osteoblast markers.

#### SnRNA-Seq Analysis Confirms the Mesenchymal-Like Fate Transition of Cardiomyocytes in *Hgs* Knockout Mice

To further validate and characterize the transition of cardiomyocytes, we performed snRNA-seq analysis on heart samples from control (*α-MHC-Cre;Hgs^+/+^*, hereinafter *Hgs^+/+^*) and *Hgs* knockout (*α-MHC-Cre;Hgs^fl/fl^*, hereinafter *Hgs^fl/fl^*) mice. We profiled a total of 20,120 nuclei from both groups. Unsupervised clustering via a uniform manifold approximation and projection (UMAP) identified 6 major cell populations, which were further resolved into 25 transcriptionally distinct subclusters. These included 4 cardiomyocyte, 8 fibroblast, 7 macrophage, 2 endocardial, 3 endothelial and 1 mural cell subclusters (Figure S6A). Signature genes defining each cluster are shown in a heatmap (Figure S6B). Consistent with histological observations, *Hgs^fl/fl^* hearts exhibited increased fibroblast proportions (Figure S6C), confirming the fibrotic phenotype at the transcriptomic level.

GO pathway enrichment analysis revealed that downregulated genes in *Hgs* knockout cardiomyocytes were primarily involved in muscle development, mitochondrial organization, and heart contraction—hallmarks of cardiomyocyte function (Figure S7A). Conversely, upregulated genes were associated with ECM organization, mesenchyme development, collagen biosynthesis, and ossification, indicating a transition towards mesenchymal-like cells (Figure S7A). Differential expression analysis further confirmed this transition, showing decreased expression of cardiomyocyte identity markers (*Myh6*, *Tnnt2*, *Atp2a2, Ryr2*) alongside increased mesenchymal transcription factors (*Twist1*, *Snail*), mesenchymal cell markers (*Vim*, *Pdgfrα*), myofibroblast marker *α-SMA*, ECM-associated molecules (*Postn*, *Fap*, *Fn1*, *Col1a2, Pcolce*, *Vcan*), and osteogenic markers (*Runx2*, *Spp1*, *Sparc*, *Enpp1*) in *Hgs* knockout cardiomyocytes (Figure S7B).

We next focused on the 4 cardiomyocyte subclusters (CM1–CM4), each exhibiting distinct transcriptional profiles: CM1 was enriched genes encoding ion-channels (*Kcnj3*, *Kccn2*, *Kcnd2*, *Grm1*) and those involved in cardiac muscle development (*Tnni3k*, *Myocd*, *Mhrt*, *Myh7b*, *Actn2*, *Atp2a2*), consistent with cardiomyocyte characteristics. CM2 showed enrichment for genes related to morphogenesis (*Shroom3*, *Cap2*) and pathological stress response (*Ankrd1*, *Rbm20*, *Col4a5*, *Tmem117*). CM3 was characterized by ECM-related genes (*Dcn*, *Col8a1*, *Col15a1*, *Fbn1*, *Mgp*), cell adhesion molecule *Pcdh9*, and mesenchymal cell marker *Pdgfra*. CM4 enriched in mitochondrial organization genes (*mt-Atp6*, *mt-Nd4*, *mt-Co2*, *mt-Cytb*), ribosome biogenesis genes (*Rplp1*, *Rps2*), ECM genes (*Col1a2*, *Col5a2*, *Fbn1*), and mesenchymal cell marker *Vim* (Figure 5A and 5B). Proportion analysis revealed a reduction in the CM1 cluster and an increase in the other 3 subclusters in *Hgs^fl/fl^* mice (Figure 5C). Consistently, we observed a progressive downregulation of mature cardiomyocyte markers and a stepwise upregulation of mesenchymal transcription factors, mesenchymal cell identity markers, ECM molecules, and osteogenic markers along the trajectory from CM1 to CM4 (Figure 5D). UMAP visualization confirmed that key mesenchymal (*Pdgfrα*, *Pdgfrβ*, *Vim*), fibrogenic (*Postn*, *Col1a2*), and osteogenic (*Runx2*, *Spp1*, *Sparc*) markers were predominantly expressed in the CM2-4 clusters of *Hgs^fl/fl^* mice (Figure S7C). This observation is consistent with an association between these transitional states and a mesenchymal–like phenotype.

**Figure 5.**
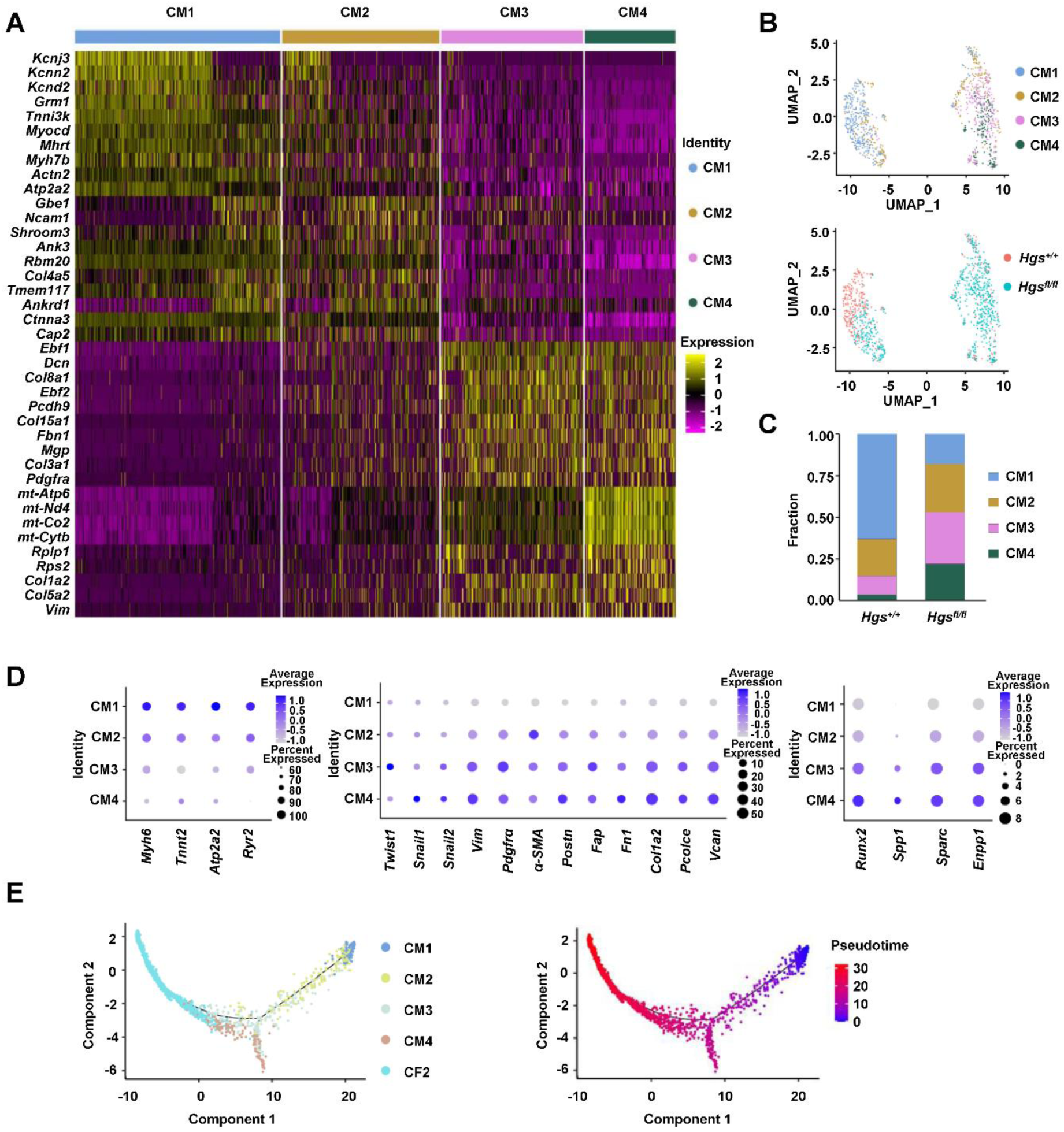
SnRNA-Seq analysis confirms the mesenchymal-like fate transition of cardiomyocytes in *Hgs* knockout mice. **A**, Heatmap displaying distinct gene expression profiles among cardiomyocyte subclusters. **B**, UMAP projection identifying cardiomyocyte subtypes in hearts of mice. **C**, Proportional analysis of each cardiomyocyte subcluster in hearts from mice. **D**, Dot plots showing expression levels of representative marker genes for cardiomyocytes, mesenchymal cells, activated fibroblasts, and osteoblasts in each cardiomyocyte subcluster (CM1 to CM4). **E**, Pseudotime analysis mapping the transition trajectory from cardiomyocytes to mesenchymal cells.

To infer potential lineage relationships, we performed pseudotime trajectory analysis (Monocle version 2.26.0) integrating cardiomyocytes and the adjacent mesenchymal CF2 cluster (Figure S6A). The resulting trajectory revealed a continuous progression from CM1 through CM2, CM3, and CM4 toward the CF2 mesenchymal cluster, providing computational evidence suggestive of a mesenchymal-like fate transition of cardiomyocytes in *Hgs^fl/fl^* mice (Figure 5E).

Altogether, these data confirm that *Hgs* deletion in cardiomyocytes induces a progressive mesenchymal-like fate transition, underscoring the role of HGS in maintaining cardiomyocyte identity.

#### *Aldh1a2* is Upregulated in Cardiomyocytes of Fibrotic Hearts

To investigate the mechanism driving the mesenchymal-like fate transition upon *Hgs* deletion, we integrated proteomic data^49^ and snRNA-seq data from *Hgs* knockout mice. Proteomic analysis identified top 6 EMT-associated proteins^50–55^ significantly upregulated in *Hgs*-deficient hearts, including CREB1, NQO1, STRATIFIN, PIRIN, RGS10, and ALDH1A2 (Figure 6A). ALDH1A2, a critical rate-limiting enzyme in retinoic acid signaling, was selected for further study based on its established role in fibroblast-to-myofibroblast transition^56^ and its upregulation in cardiomyocytes of *Hgs^fl/fl^* mice, particularly in the CM4 cluster in our snRNA-seq data (Figure 6B).

**Figure 6.**
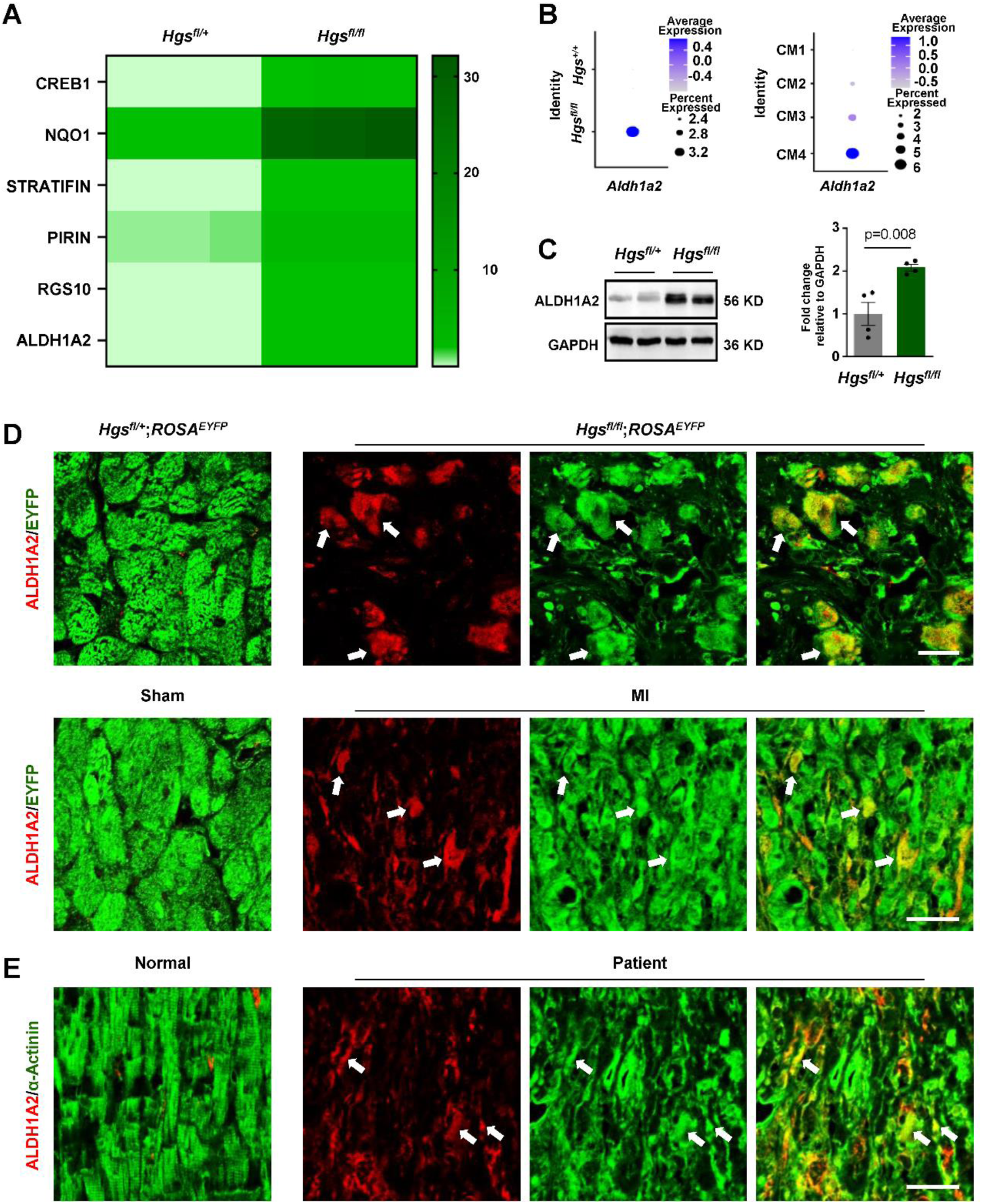
*Aldh1a2* is upregulated in cardiomyocytes of fibrotic hearts. **A**, Heatmap of the top upregulated EMT-associated proteins identified in proteomic data. **B**, Dot plots illustrating *Aldh1a2* expression levels in cardiomyocytes from mice and in each cardiomyocyte subcluster. Color intensity indicates expression levels. Dot size reflects the percentage of cells expressing *Aldh1a2*. **C**, Western blot analysis of ALDH1A2 protein levels in hearts from mice, with quantification shown to the right (n = 4). Data were presented as means ± SEM. *P* values were determined by unpaired two-tailed Student’s *t* test. **D**, Representative immunofluorescence images of co-expression of ALDH1A2 (red) with EYFP (green) in heart sections from mice. Scale bars, 25 μm. **E**, Representative immunofluorescence images of co-expression of ALDH1A2 (red) with EYFP (green) in heart sections from human. Scale bar, 25 μm.

We confirmed significant upregulation of ALDH1A2 in the hearts of *Hgs* knockout mice by western blot analysis (Figure 6C). Immunofluorescence staining further demonstrated that a subset of cardiomyocyte-derived EYFP^+^ cells co-expressed ALDH1A2 in both *Hgs* knockout and MI mice (Figure 6D). Consistent with these findings, ALDH1A2 was also elevated in some cardiomyocytes from MI patients compared to those from normal human hearts (Figure 6E).

Collectively, these results indicate that ALDH1A2 is consistently upregulated in cardiomyocytes under fibrotic conditions, suggesting that it may serve as a potential mediator of their mesenchymal–like fate transition.

#### *Aldh1a2* Overexpression in Cardiomyocytes is Sufficient to Induce a Mesenchymal-Like Fate Transition

To determine whether *Aldh1a2* overexpression is sufficient to induce a mesenchymal-like fate transition, we transduced primary neonatal cardiomyocytes with an adenovirus encoding *Aldh1a2* (*Ad-Aldh1a2*) (Figure 7A). RT-qPCR analysis showed that *Ad-Aldh1a2* transduction significantly upregulated the expression of mesenchymal marker genes, including *α-SMA*, *Postn*, and *Col1a1*, compared to controls (Figure 7B).

**Figure 7.**
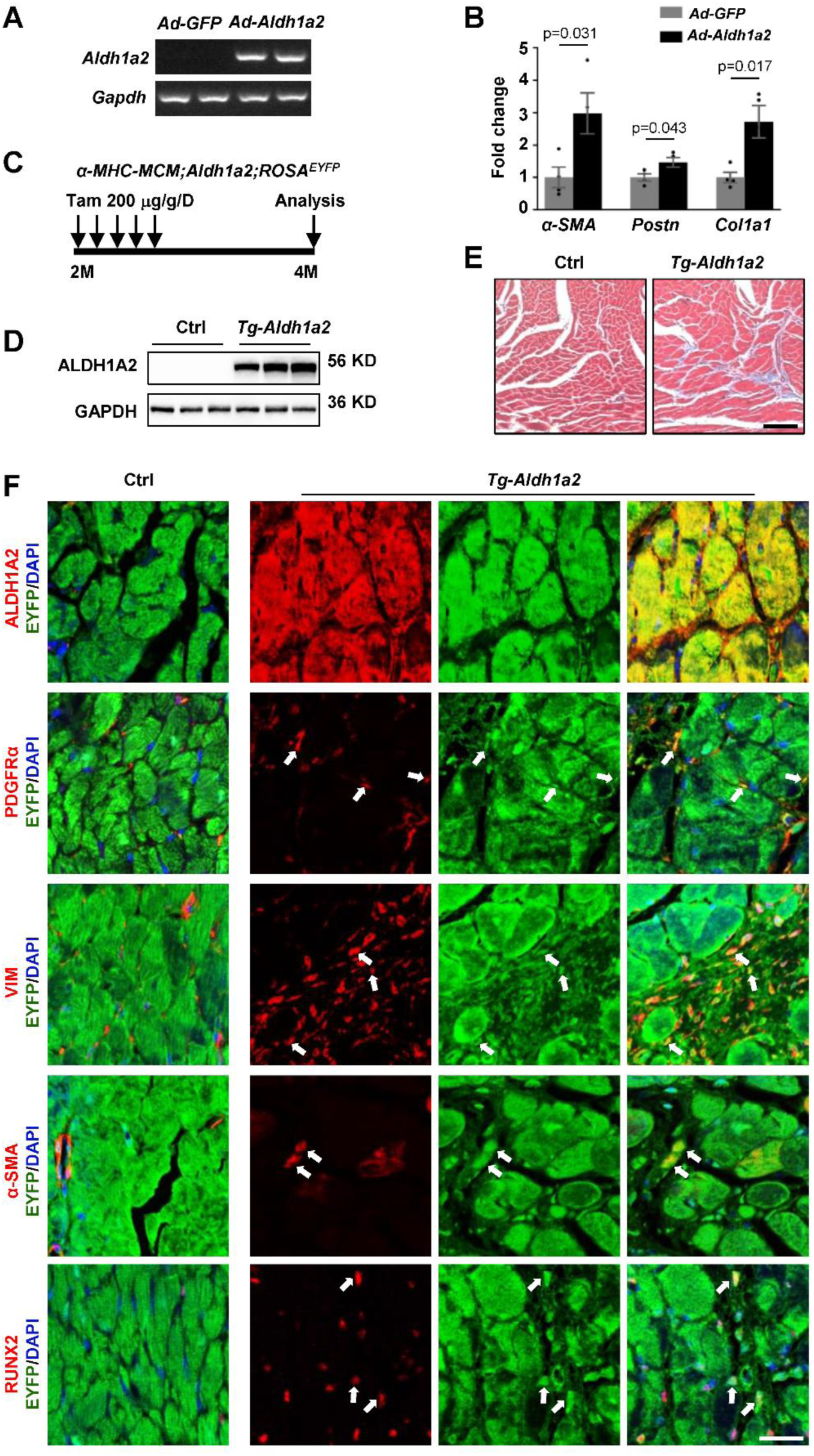
*Aldh1a2* overexpression in cardiomyocytes is sufficient to induce a mesenchymal-like fate transition. **A**, RT-PCR analysis showing *Aldh1a2* mRNA levels in primary neonatal cardiomyocytes treated with Ad-GFP or Ad-Aldh1a2. **B**, RT-qPCR analysis of mesenchymal marker gene expression in primary neonatal cardiomyocytes treated with Ad-GFP or Ad-Aldh1a2 (n = 4). Data were presented as means ± SEM. *P* values were determined by unpaired two-tailed Student’s *t* test. **C**, Schematic illustrating the lineage tracing study design in *α-MHC-MCM;Aldh1a2;ROSA^EYFP^* (*Tg-Aldh1a2*) mice. **D**, Western blot analysis of ALDH1A2 protein levels in *α-MHC-MCM;ROSA^EYFP^* (Ctrl) and *Tg-Aldh1a2* hearts. **E**, Masson’s trichrome staining of cardiac sections from mice. Scale bar, 100 μm. **F**, Representative immunofluorescence images showing co-expression of ALDH1A2 (red) or mesenchymal markers (red) with EYFP (green) in heart sections from mice. Scale bar, 25 μm.

To validate the functional role of *Aldh1a2 in vivo*, we generated an inducible cardiomyocyte-specific *Aldh1a2* overexpression mouse line (*α-MHC-MCM;Aldh1a2;ROSA26^EYFP^*, hereinafter *Tg-Aldh1a2*). In this system, *Aldh1a2* overexpression in EYFP-labeled cardiomyocytes and their progeny was triggered by tamoxifen treatment (Figure S8). *Tg–Aldh1a2* mice and their littermate controls (*α-MHC-MCM*;*ROSA26^EYFP^*) were treated with tamoxifen at 2 months of age and analyzed 2 months post–induction (Figure 7C). 4–month–old *Tg–Aldh1a2* mice exhibited robust ALDH1A2 protein overexpression in cardiomyocytes, as confirmed by western blot and immunofluorescence staining (Figure 7D and 7F).

Notably, these *Tg–Aldh1a2* mice developed myocardial fibrosis, evidenced by extensive collagen deposition in Masson’s trichrome–stained heart sections (Figure 7E). Lineage–traced immunofluorescence analysis showed that a subset of EYFP^+^ cardiomyocyte–derived cells in *Tg-Aldh1a2* mice co-expressed the mesenchymal markers PDGFRα, VIM, α-SMA, and RUNX2, suggesting that ALDH1A2 overexpression *in vivo* is sufficient to induce a transition toward mesenchymal-like cells (Figure 7F).

Together, these results suggest that enforced expression of *Aldh1a2* is sufficient to drive a mesenchymal-like fate transition and contribute to fibrotic remodeling, supporting a functional role for ALDH1A2 in cardiomyocyte plasticity during myocardial fibrosis.

#### *Aldh1a2* Knockout in Cardiomyocytes Attenuates MI-Induced Myocardial Fibrosis

To assess the role of *Aldh1a2* in MI-induced myocardial fibrosis, we generated cardiomyocyte-specific *Aldh1a2* knockout (*α-MHC-Cre;Aldh1a2^fl/fl^*, hereinafter *Aldh1a2^fl/fl^*) mice. 8-week-old *Aldh1a2^fl/fl^* mice and their littermate controls were subjected to sham or MI operation and analyzed after 3 weeks. MI induced a marked upregulation of ALDH1A2 in control mice, which was significantly attenuated in *Aldh1a2^fl/fl^* hearts (Figure 8A). Masson’s trichrome staining revealed extensive myocardial fibrosis in control mice post-MI, whereas fibrosis was attenuated in *Aldh1a2^fl/fl^* hearts (Figure 8B and 8C). Besides, MI-induced elevated expression of fibrosis-related genes (*Col1a1* and *Col3a1*) and cardiac remodeling markers (*Nppa*, *Nppb*, and *Acta1*) was significantly decreased in *Aldh1a2^fl/fl^* hearts (Figure 8D and 8E). Echocardiograph analysis revealed that MI-induced increases in left ventricular diastolic internal diameter (LVID;d), volume (LV VOL;d), and mass (LVM), were all significantly attenuated in *Aldh1a2^fl/fl^* mice (Figure 8F; Table S2). Additionally, the MI-induced decrease in ejection fraction (EF) and fractional shortening (FS) showed a trend toward recovery in these mice, albeit without statistical significance (Figure 8F; Table S2).

**Figure 8.**
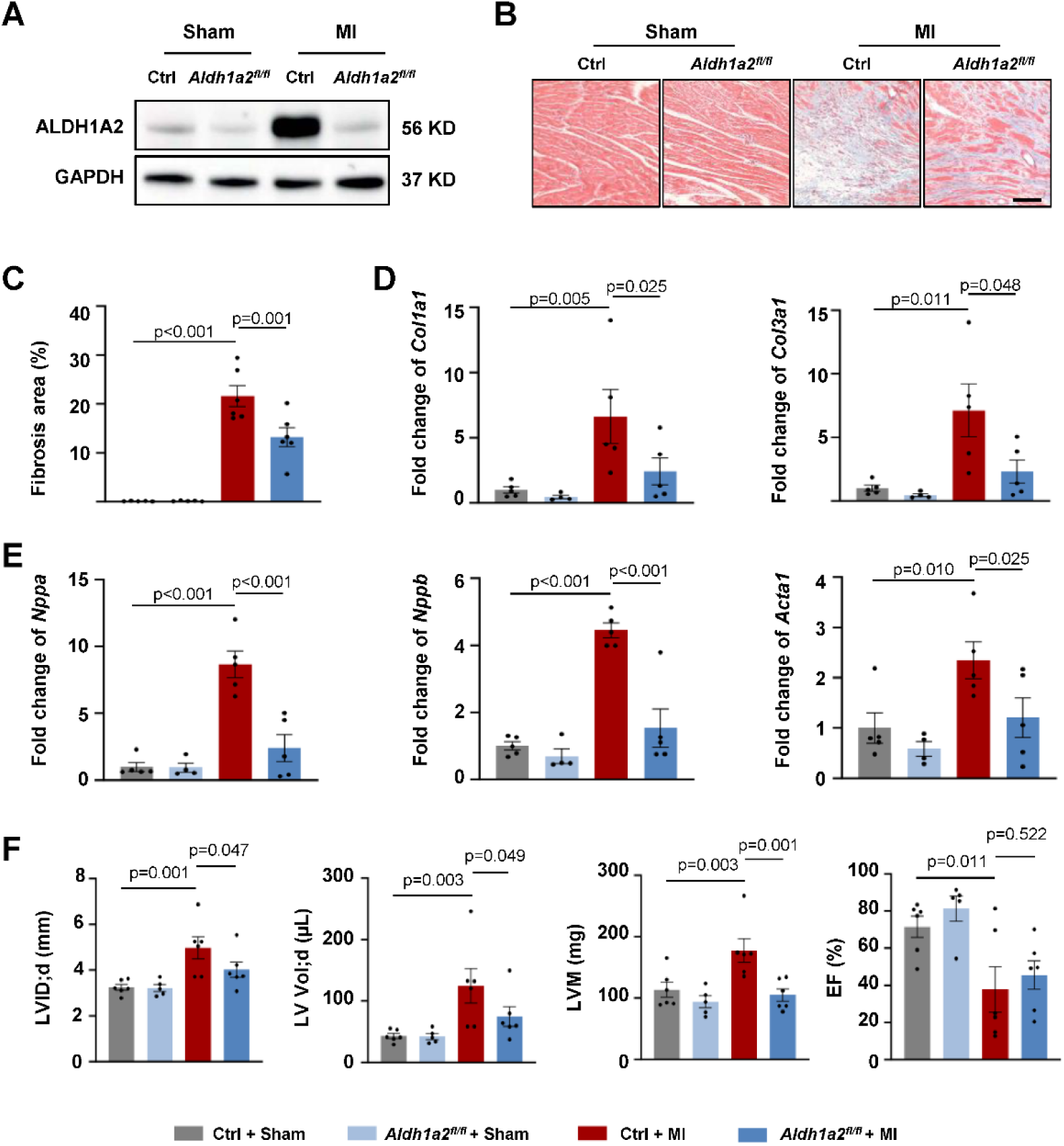
Knockout of *Aldh1a2* in cardiomyocytes attenuates MI-induced myocardial fibrosis. **A**, Western blot analysis of ALDH1A2 expression in mice. **B**, Masson’s trichrome staining of hearts from mice. Scale bar, 100 µm. **C**, Quantification of the fibrotic areas of hearts from mice. **D** and **E**, RT-qPCR analysis of fibrosis and pathological remodeling marker genes in mouse heart tissues. **F**, Measurement of LV internal dimension of in diastole (LVID;d), LV volume in diastole (LV Vol;d), LV mass (LVM), and ejection fraction (EF) in mice. Data in **C**-**F** were presented as mean ± SEM, *P* value was calculated by two-way ANOVA followed by Tukey’s multiple comparisons test. Number of samples is shown on bar graphs.

These findings demonstrate that cardiomyocyte-specific deletion of *Aldh1a2* attenuates MI-induced myocardial fibrosis, highlighting ALDH1A2 as a potential therapeutic target for pathological cardiac remodeling.

## DISCUSSION

The plasticity of adult cardiomyocytes has remained a long-standing subject of debate. Here, we provide *in vivo* genetic evidence that, in the setting of myocardial fibrosis, a subset of adult cardiomyocytes can undergo a mesenchymal-like fate transition and subsequently give rise to cells expressing markers of fibroblasts or osteoblasts. Mechanistically, we demonstrate that *Hgs* deficiency derepresses *Aldh1a2*, and that either loss of *Hgs* or transgenic *Aldh1a2* overexpression is sufficient to drive this maladaptive cellular transition. Conversely, cardiomyocyte-specific *Aldh1a2* knockout attenuates MI-induced myocardial fibrosis.

Our findings support the concept that adult cardiomyocytes, under sustained pathological stress, can undergo a mesenchymal-like fate transition. These observations align with and extend the emerging concept of mesenchymal drift—loss of original cellular identity and acquisition of mesenchymal traits—recently described across multiple cell types during aging and disease by Lu et al.^57^ In the cardiac context, we provide genetic evidence that this transition is characterized by the coordinated downregulation of cardiomyocyte identity markers and the activation of a mesenchymal program. Key features include upregulation of mesenchymal transcription factors, expression of fibroblast and osteoblast markers, acquisition of a spindle-shaped morphology with disrupted cell–cell contacts and loss of alignment, expression of proliferation markers, and transcriptional enrichment for genes related to ECM organization, migration, and fate determination. This finding broadens the spectrum of adult cardiac plasticity beyond the well-described EMT or EndMT observed during cardiac development and regeneration.^7^ While cardiomyocyte dedifferentiation, proliferation and redifferentiation have been documented during mammalian heart regeneration,^25,29,58,59^ whether adult cardiomyocytes retain the capacity for a mesenchymal-like fate transition in adult fibrotic disease has remained unclear. Through integrated analysis of human MI specimens and rigorous genetic lineage tracing in murine models of MI and *Hgs* knockout, we provide evidence that a subset of cardiomyocytes indeed undergoes such a transition. Importantly, dual recombinase-mediated lineage tracing provides genetic evidence that some cardiomyocyte-derived cells not only undergo a mesenchymal-like fate transition but also give rise to cells expressing myofibroblast or osteoblast markers. Although this phenotypic transition may represent an adaptive survival response under pathological stress, its net impact on long-term cardiac function warrants further investigation. Our findings align with emerging evidence that diverse muscle cell types can undergo mesenchymal transitions under specific conditions,^60,61^ suggesting that such phenotypic adaptability may be an evolutionarily conserved feature of muscle tissues.

We identify HGS and ALDH1A2 as regulators of the mesenchymal-like potential of adult cardiomyocytes. Genetic deletion of *Hgs* derepresses this potential, resulting in a mesenchymal-like fate transition accompanied by myocardial fibrosis and calcification. Further investigation reveals ALDH1A2 as a downstream effector driving this pathogenic transition. Although ALDH1A2 has been previously implicated in epicardial EMT, fibroblast activation, cardiac development, and injury responses in the heart,^55,56,62,63^ and its inhibition has been associated with a mesenchymal-like phenotype in tumor cells,^64^ its specific role in governing the cellular identity of adult cardiomyocytes has remained largely unknown. Here, we show that ALDH1A2 expression is significantly upregulated in cardiomyocytes of MI patients and mice, as well as in *Hgs* knockout mice. Importantly, transgenic overexpression of *Aldh1a2* in cardiomyocytes not only induces the expression of mesenchymal and myofibroblast markers but also promotes overt myocardial fibrosis, whereas *Aldh1a2* knockout in cardiomyocytes partly alleviates MI-Induced myocardial fibrosis. These findings may appear to differ from a prior report demonstrating that cardiomyocyte ALDH1A2 exerts cardioprotective effects in acute I/R injury.^65^ This apparent discrepancy may reflect the context-dependent pleiotropic nature of RA signaling: transient ALDH1A2 upregulation in acute injury may engage cytoprotective programs, whereas sustained or supraphysiological activation—as modeled here—may shift RA signaling output toward mesenchymal reprogramming. Collectively, these findings identify ALDH1A2 as a regulator of the mesenchymal-like fate transition in cardiomyocytes and reveal a mechanistic link between this pathway and fibrotic heart disease, suggesting its potential as a therapeutic target.

This study establishes cardiomyocytes as a previously underappreciated cellular source of cardiac fibroblasts during myocardial fibrosis. Although resident fibroblasts derived from embryonic epicardium or endothelial cells constitute the major source of activated fibroblasts in the adult heart,^7,14–16^ with additional contributions from other cardiac cell types, including pericytes, monocytes, CD34^+^ cells, and macrophages,^21–23,66^ our findings indicate that a fraction of cardiomyocytes themselves can undergo a mesenchymal-like fate transition and give rise to cells expressing fibroblast or myofibroblast markers. The expression of mesenchymal transcription factors and markers in cardiomyocytes from MI patients corroborates previous observations of mesenchymal marker expression in pressure overload–induced cardiac hypertrophy mouse model,^37^ suggesting that this plasticity may reflect a general response to biomechanical stress. Genetic lineage tracing confirms that cardiomyocyte-derived mesenchymal-like cells contribute to the pool of myofibroblasts or osteoblasts in MI and *Hgs* knockout mice. Furthermore, RNA-seq analysis revealed that, compared with mesenchymal cells from other origins, cardiomyocyte–derived mesenchymal–like cells exhibit a highly active transcriptional signature characterized by elevated expression of genes involved in cell adhesion, ECM organization, collagen production, migration, and proliferation, together with distinct enrichments in genes related to cell morphology, fate determination, and osteogenesis. This profile suggests an enhanced matrix-synthetic and migratory capacity that could actively contribute to fibrotic remodeling. These findings expand our understanding of cellular plasticity and the diversity of fibroblast origins in cardiac disease.

Several important questions remain. First, although we demonstrate that some cardiomyocytes can acquire mesenchymal characteristics and contribute to the fibroblast pool, the quantitative contribution of these cells to the total fibrosis burden and their functional impact needs to be precisely determined. Second, the molecular circuitry that initiates and sustains this fate transition remains incompletely understood. Third, the fragility and low abundance of these transitioned cells have precluded the application of conventional functional assays. Addressing these gaps will be essential for translating these insights into targeted anti-fibrotic therapies that preserve cardiac function.

## ARTICLE INFORMATION

### Affiliations

State Key Laboratory of Medical Proteomics, National Center for Protein Sciences (Beijing), Beijing, China (T.W., M.L., Y.X., C.H., Z.L., Y.T., G.Y., W.L., P.X., J.W., X.Y.); Peking-Tsinghua Center for Life Sciences, Academy for Advanced Interdisciplinary Studies, Center for Quantitative Biology (CQB), Peking University, Beijing 100871, China (C.Z., R.L., J.H.); School of Life Sciences, Hebei University, Baoding, Hebei, China (M.L., Y.X.); State Key Laboratory of Membrane Biology, College of Life Sciences, Peking University, Beijing, China (Y.H., S.W.); New Cornerstone Science Laboratory, Key Laboratory of Multi-Cell Systems, Shanghai Institute of Biochemistry and Cell Biology, Center for Excellence in Molecular Cell Science, Chinese Academy of Sciences, University of Chinese Academy of Sciences, Shanghai, China (B.Z.).

## Acknowledgements

We thank Dr. Jiangping Song from Fuwai Hospital for providing heart samples of normal human and MI patients.

## Supplemental Material

Supplemental Methods

Figures S1–S8

Tables S1–S2

## NONSTANDARD ABBREVIATIONS AND ACRONYMS

α-SMA: α-smooth muscle actin
Aldh1a2: aldehyde dehydrogenase 1 family member A2
cTnT: cardiac troponin T
ECM: extracellular matrix
EMT: epithelial-mesenchymal transition
EndMT: endothelial-mesenchymal transition
ESCRT: endosomal sorting complex required for transport
GO: gene ontology
Hgs: hepatocyte growth factor-regulated tyrosine kinase substrate
MI: myocardial infarction
PDGF: platelet-derived growth factor
POSTN: periostin
snRNA-seq: single-nucleus RNA sequencing
TGF-β: transforming growth factor β
UMAP: uniform manifold approximation and projection
VIM: vimentin

